# Cost-effective early detection of banana bunchy top disease: insights from spatio-temporal modelling in Benin

**DOI:** 10.64898/2026.01.09.698707

**Authors:** Renata Retkute, Christopher A. Gilligan

## Abstract

Banana bunchy top disease (BBTD), caused by banana bunchy top virus (BBTV), threatens food security and livelihoods across sub-Saharan Africa. Effective surveillance is critical for early detection, but in many regions, monitoring is constrained to short operational windows due to limited funding and laboratory capacity. When surveillance is restricted to a single calendar year, sampling effort must be carefully allocated to maximize early detection, yet quantitative guidance linking detection performance to cost is lacking. To address this, we simulated the spatio-temporal spread of BBTV in Benin and retrospectively evaluated country-level BBTD surveillance using a single-year cross-sectional survey design. We found that early detection is feasible but prohibitively expensive (USD 100,000 per year). Under a constrained budget of USD 10,000 per year, an optimal strategy - defined as reaching 75% mean detection probability earliest - was 500 sites with 10 samples per site, though this could delay BBTD detection by up to one year. Our integrated simulation - economic framework quantifies trade-offs between cost and detection likelihood, providing guidance for resource-efficient national-scale BBTD surveillance and a transferable approach for plant pathogen monitoring in smallholder systems.

## 1. Introduction

Banana and plantain (*Musa spp.*) are among the most important food crops in sub-Saharan Africa, underpinning household nutrition and generating income for millions of smallholder farmers. Across diverse agroecological zones, banana production supports both subsistence consumption and local markets, making it a cornerstone of rural livelihoods and food systems [30]. However, sustained productivity is increasingly undermined by biotic constraints, particularly viral diseases. Banana bunchy top disease (BBTD), caused by banana bunchy top virus (BBTV; *Babuvirus musae*), is widely regarded as one of the most destructive banana diseases worldwide [20]. Transmission occurs primarily through the banana aphid, *Pentalonia nigronervosa*, which spreads the virus in a persistent, circulative, non-propagative manner following extended feeding on infected plants [26]. Beyond vector-mediated transmission, the movement of infected vegetative planting material—particularly suckers exchanged through informal farmer networks—plays a critical role in long-distance spread [29]. Infected plants exhibit distinctive symptoms, including discontinuous dark green streaks on leaf tissues, progressively narrow and brittle emerging leaves, and a characteristic “bunchy” appearance due to shortened internodes. These symptoms often culminate in severe stunting and failure to produce a harvestable bunch, resulting in substantial yield losses or total crop failure if the disease is not controlled [14, 28].

Banana bunchy top virus has a long history in eastern Africa, where it occurs widely across multiple production landscapes and is recognised as a major constraint on banana and plantain cultivation, with confirmed presence in countries such as Uganda, Rwanda, Burundi and neighbouring regions within the Democratic Republic of Congo (DRC) and other central African production zones [13]. In contrast, BBTV emergence in West Africa is relatively recent, with the first confirmed reports occurring only in the past decade. The virus was initially detected in Benin in 2011, where molecular diagnostics confirmed BBTV infection in symptomatic *Musa* plants [15]. This was followed by confirmation in Nigeria in 2012 [1], and later in Togo in 2018 [12].

Surveillance of plant pathogens is essential for national and regional plant health systems, providing early warning of pests that threaten food security, trade, and the environment, and enabling rapid response, informed phytosanitary decisions, and effective risk management [18]. International guidance from the FAO and IPPC’s Africa Phytosanitary Programme provides practical protocols for BBTV field surveys, including standardized sample collection procedures that maximize viral titre while minimizing the risk of cross-contamination [27]. Despite the existence of international guidance, there are no quantitative studies explicitly linking BBTV surveillance performance to economic cost.

In practice, plant health surveillance is often constrained by limited funding and laboratory capacity, supporting monitoring activities over short operational windows rather than sustained annual programmes. When surveillance can be conducted in only a single calendar year, decision-makers must carefully allocate sampling effort across space and time to maximise the probability of early pathogen detection. Retrospective analyses of simulated epidemics provide a powerful framework to quantify how sampling intensity, spatial coverage, and underlying prevalence jointly influence detection probability and surveillance cost. Such model-based approaches have been widely applied in animal and human health, including optimisation of SARS-CoV-2 detection strategies [19], evaluation of facility-level malaria surveillance sensitivity [4], early-warning monitoring for foot-and-mouth disease incursions [7], and assessment of bovine tuberculosis surveillance in badger populations [11].

An expanding literature on modelling research has examined how plant disease surveillance can be optimised prior to outbreaks, typically using prospective frameworks in which hypothetical pathogen incursions are simulated and alternative sampling designs are evaluated [17]. For example, a spatially explicit optimisation approach integrating epidemiological spread of citrus disease huanglongbing with surveillance design was developed in [16], demonstrating that targeting only the highest-risk sites is often a suboptimal strategy for early detection once spatial dynamics are taken into account. In a sub-Saharan African context, integrated landscapeand fieldscale simulations were used to assess surveillance strategies for the early detection of cassava brown streak disease under assumed introductions in Nigeria, showing that allocating survey effort in proportion to host density improves detection relative to uniform sampling [9].

In this study, we retrospectively evaluated the performance and cost of country-level BBTD surveillance conducted within a single calendar year. We first simulated the spatio-temporal spread of BBTV across Benin using host maps and epidemiological parameters estimated by Retkute et al. [25] and then used these epidemics to evaluate alternative cross-sectional surveillance designs. By integrating epidemic simulations with an explicit cost framework, we quantified trade-offs between detection likelihood and surveillance expenditure across a range of sampling strategies involving different numbers of surveyed sites and numbers of sampled plants per site. The objective is to provide evidence-based guidance for designing resource-efficient BBTV surveillance programs for first detection of an incursion under realistic budgetary constraints.

## 2. Methods

### 2.1. Modelling BBTD spread

Historical spread of BBTD across Benin was simulated using the banana production map at 250m resolution and the model developed in Retkute et al. [25]. The BBTV spread model uses a spatially explicit susceptible–infected (S–I) framework that accounts for variations in banana density, multiple pathogen introduction pathways, localized increases in pathogen load, and virus transmission between grid cells. Each cell is characterized by the local banana host density, representing the proportion of land under banana cultivation. Grid cells are assigned to one of two epidemiological states: susceptible (S) or infected (I). A cell is considered susceptible if all banana plants within it are healthy, and transitions to the infected state once the first plant becomes infected. Infection of a susceptible cell can occur via primary transmission, representing introductions from external sources (e.g., long-distance human-mediated movement), or via secondary transmission from infectious neighbouring cells, typically mediated by the banana aphid vector [24]. Following the initial infection event, disease progression within an infected cell is governed by a deterministic local transmission process [23]. Full model description and parameterisation are provided in [25].

### 2.2. BBTD occurrence data

Banana bunchy top virus was first detected in Benin in 2011 [15], and these initial infection sites were used to seed the simulated epidemic for that year. To evaluate how well the model reflected observed BBTD spread, we compared simulated commune-level timings of first infection with the limited available empirical data. Two independent surveillance datasets, published after the initial incursion, were used for comparison: occurrence records from the 2017 regional survey [31] and field assessments from 2019 [5]. As surveillance data from [25] were used for model parameterisation, they were excluded from the validation analysis.

### 2.3. Operational and economic assumptions for BBTD surveillance implementation

In this study, surveillance was modelled as a cross-sectional survey conducted within one calendar year, rather than as a recurring annual program. The probability of detecting at least one infected plant was evaluated under a country-level sampling framework (i.e. to simulate first detection of the disease). At the plant level, we assumed perfect diagnostic sensitivity and specificity.

Sampling units (grid cells) were chosen using simple random sampling, whereby each eligible grid cell in the study landscape had an equal probability of being included in the surveillance sample, in the absence of any assumed prior knowledge about likely sites for initial infection. For a set of *N* surveillance sites and *K* samples collected per site, with each site having specific prevalence of BBTD, *p_i_*, the probability of detection was defined as:

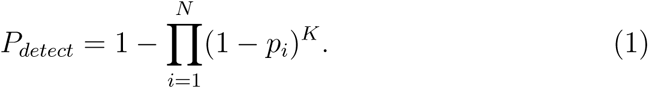

We assumed that a single surveyor could fully survey up to 10 sites per day, inclusive of travel time between sites. For analytical tractability, we assumed that the time required to collect samples within a site was negligible relative to inter-site travel and therefore did not vary with the number of samples collected per site (e.g., 1 versus 50 samples).

For context in Benin, statutory labour market data indicate that the national minimum wage is approximately XOF 52,000 per month ( USD 84), equating to roughly USD 3–4 per working day under a standard 40- hour week [21]. We adopted a conservative daily surveillance cost of USD 10 per surveyor, which encompasses field labour, transport allowances, and an allocation for equipment depreciation in line with expenditure categories used in FAO surveillance cost analyses [10].

Commercial ELISA reagent kits are available for the serological detection of BBTV and are typically sold in multi-sample formats (e.g., 96, 500, 1000, 50000 well kits) [3]. These assays provide 100% diagnostic sensitivity and 100% diagnostic specificity for samples collected from banana leaves [2]. General research-grade ELISA kits for viral antigen or antibody detection in plant pathology are broadly priced in the range of USD 300–800 per 96-well kit, suggesting reagent costs 3.1 to 8.3 USD per sample [6]. For the purposes of our analysis, we assumed that costs could be negotiated toward the lower end, resulting in a per-sample reagent cost of 3 USD.

Based on these assumptions, the total cost of a surveillance program was calculated as:

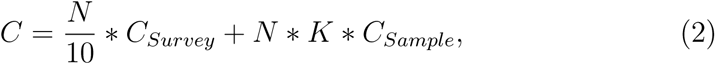

where *C_Survey_*is the daily cost per surveyor, *C_Sample_* is the reagent cost per sample, *N* is the number of sites surveyed, and *K* is the number of samples collected per site.

## 3. Results

### 3.1. Reconstructed spatio-temporal spread of BBTD in Benin

Simulations reconstructed a coherent spatio-temporal pattern of progressive spatial spread of BBTD in Benin from 2011 to 2025 (Fig. 1). Simulated epidemics exhibited an initial lag phase, during which spatial expansion was limited (2011-13; not shown). This was followed by a sustained phase of epidemic growth characterized by progressive spatial spread evident during 2014, with a gradual northward expansion of the epidemic. Early spread was concentrated in the southern and south-western departments, particularly Mono, Couffo, and Atlantique, before extending eastward and northward into Zou, Ouémé, Plateau, Collines, and subsequently the northern departments.

**Figure 1:**
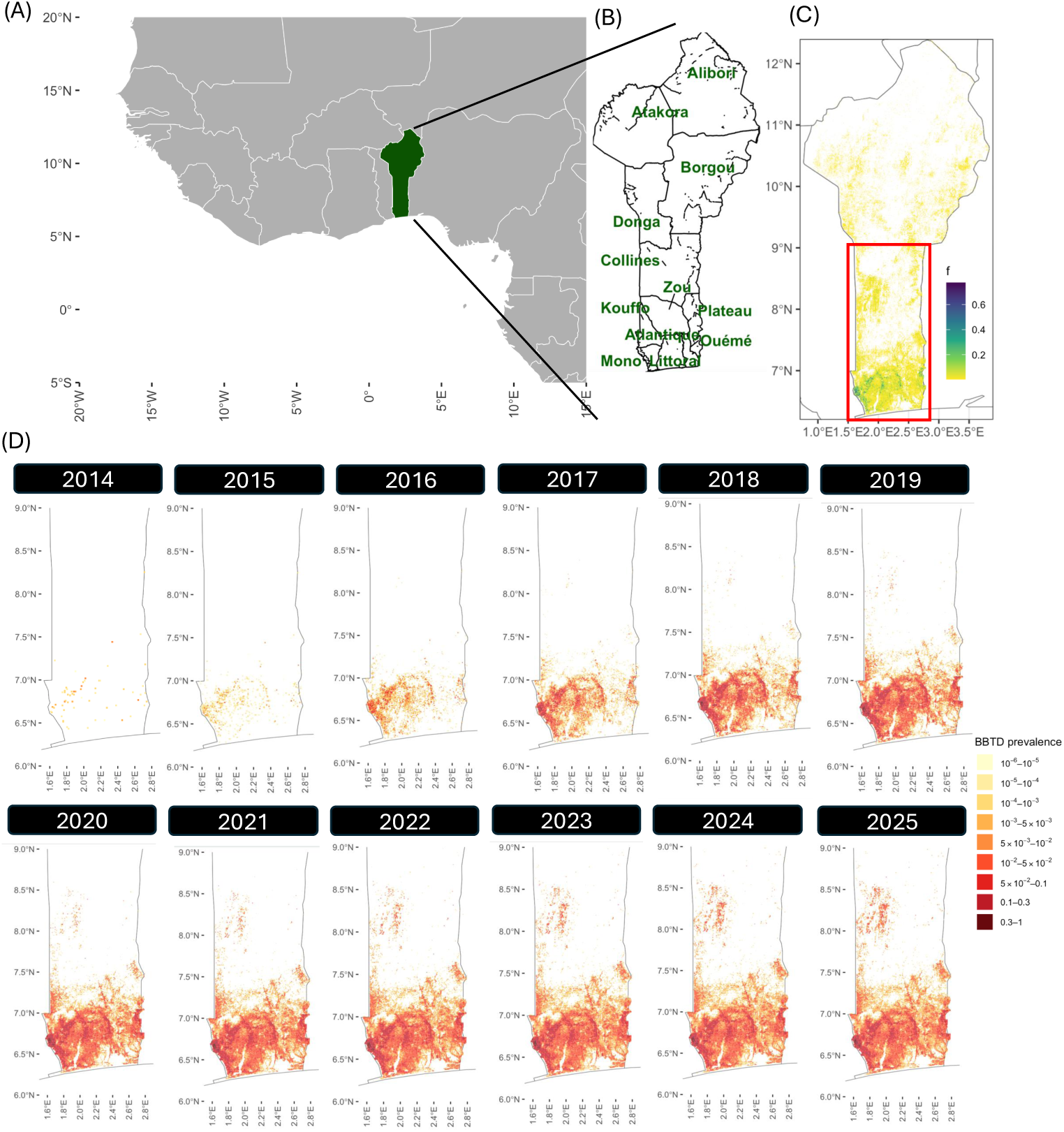
Reconstructed spread of banana bunchy top disease (BBTD) in Benin.(A) Location of Benin in West Africa. (B) Departments in Benin. (C) Distribution map for banana cultivation in Benin (produced from [25]). Shading shows fraction of a 250m × 250m grid cell planted with bananas. Red square indicates region shown in D. (D) Example simulation showing the spatial evolution of BBTD prevalence across Benin from 2014 to 2025.

Across simulations, the estimated timing of first infection varied substantially among communes but followed a clear spatial gradient (Fig. 2). Communes in southern departments were consistently infected earlier, with median first infection times clustered between 2014 and 2016, whereas central and northern communes exhibited later infection times, often after 2018. Inter-quartile ranges indicated moderate uncertainty in infection timing during early expansion, increasing for communes infected later in the epidemic. Simulated timings of infection arrival aligned reasonably well with independent empirical observations. Communes where BBTD was detected in the 2017 regional survey [31] fell within the interquartile range of simulated infection times for those locations, whereas detections from surveys conducted in 2019 occurred several years after the modelled arrival of infection [5] (Fig. 1B). These comparisons suggest that the model captured the broad timing and spatial ordering of BBTD spread during the observed period, supporting its use as a basis for retrospective evaluation of surveillance performance.

**Figure 2:**
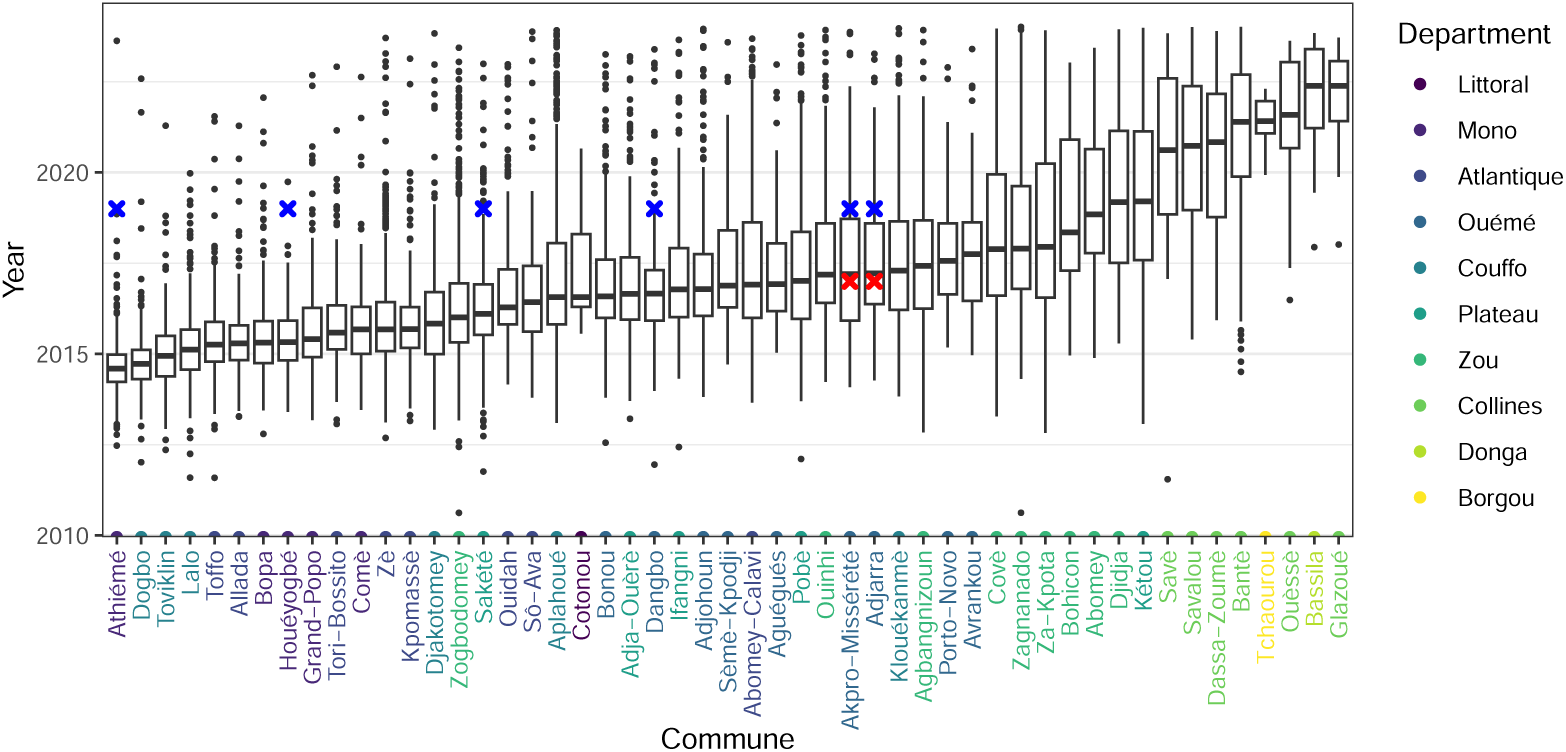
Distribution of the simulated timing of first infection at the commune level. Communes are colour-coded by department, with departments ordered longitudinally according to the centroid of each department polygon, from south (Littoral) to north (Borgou). Red crosses indicate empirical BBTD detection dates from the 2017 regional survey [31], and blue crosses indicate detection dates from field assessments conducted in 2019 [5].

### 3.2. Retrospective surveillance outcomes and cost–detection trade-offs

For the analysis, we evaluated a range of surveillance intensities by varying the number of sampled sites (*N* =10, 100, 500, and 1, 000), representing very limited to very high resource scenarios, and the number of samples collected per site (*K*= 1, 10, 20, and 50). For each surveillance configuration, we conducted 1,000 replicate simulations to quantify detection performance and associated uncertainty.

Retrospective surveillance simulations revealed strong, nonlinear interactions between survey timing, spatial coverage, and sampling intensity in determining the probability of detecting the disease (Fig. 3A). Detection performance increased with both the number of surveyed sites and the number of samples collected per site, while uncertainty declined as sampling intensity increased. Under extremely constrained resources (*N* = 10), detection probability remained low on average when a single sample was collected per site (*K* = 1), or highly uncertain when sampling intensity was increased. With moderate spatial coverage (*N* = 100), detection probabilities were negligible early in the epidemic for *K* = 1, reaching approximately 50% by 2019, whereas increasing sampling to *K* = 10 substantially accelerated detection, achieving mean probabilities exceeding 95% by 2018 but with considerable uncertainty. Further increases in within-site sampling (*K* ≥ 20) markedly reduced this uncertainty. Under high-resource scenarios (*N* = 500–1,000), near-perfect detection probabilities (*>* 95%) were achieved by 2017 for *K* ≥ 10, with expanding spatial coverage from 500 to 1,000 sites advancing reliable detection by approximately one year and substantially improving detection probabilities during the early epidemic phase.

**Figure 3:**
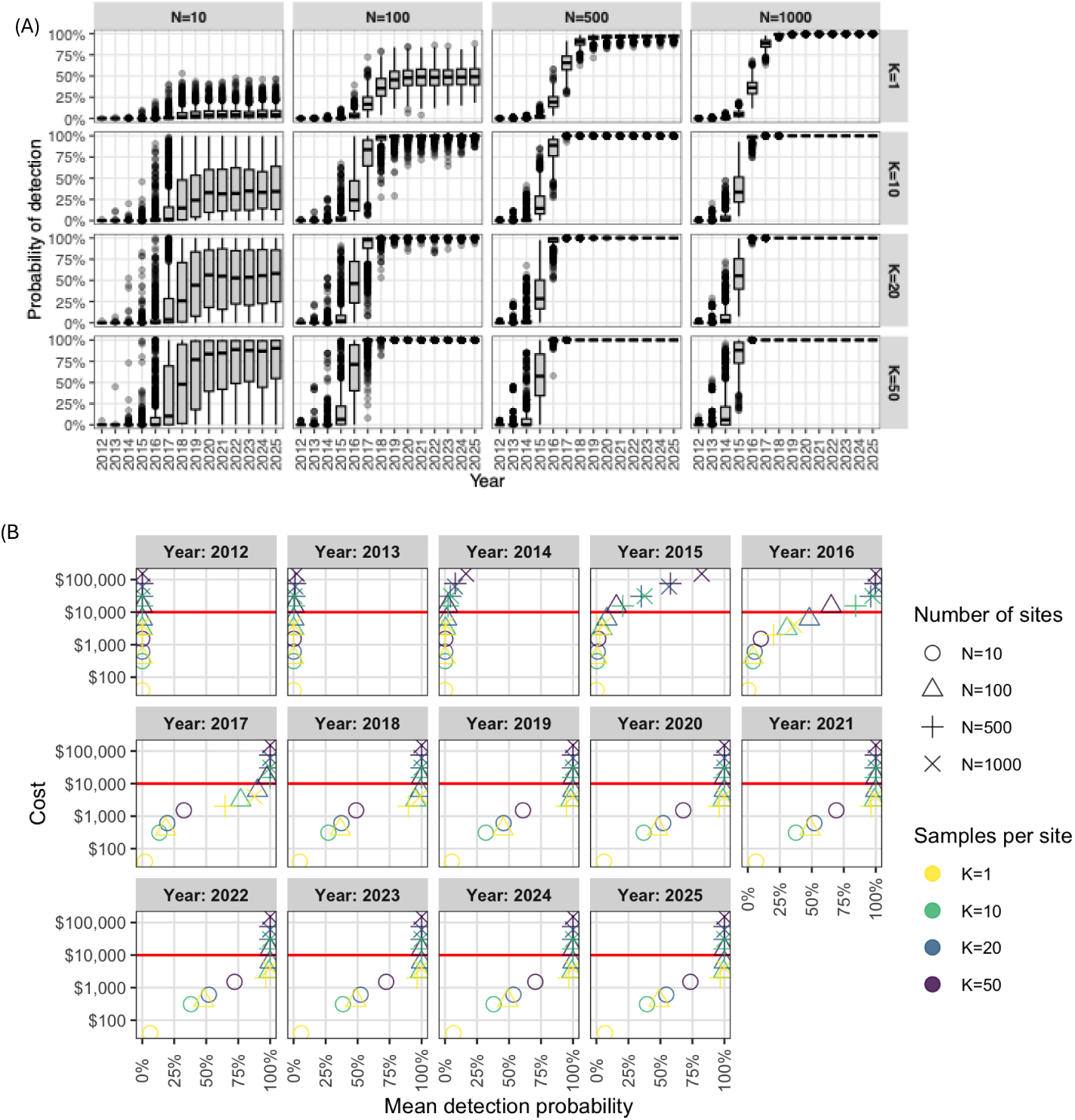
Retrospective BBTD surveillance simulations in Benin. (A) Distribution of annual detection probability over time under varying surveillance strategies: spatial coverage (*N* sites) and within-site sampling intensity (*K* samples per site). (B) Relationship between mean detection probability and total annual surveillance cost.

Cost analysis reflected a clear trade-off between detection efficiency and resource expenditure (Fig. 3B, S1). For a spatial coverage of *N* = 500 sites, total annual surveillance costs ranged from approximately USD 2,000 for minimal sampling (*K* = 1) to USD 75,500 for intensive within-site sampling (*K* = 50), while expanding spatial coverage to *N* = 1, 000 sites further amplified expenditures to USD 4,000–151,000 per year for *K* = 1−50. Temporal patterns reveal that low-intensity sampling (*K* = 1 − 10) yields very low detection probabilities (<5%) in the early epidemic years (2012–2013), which increase gradually as the epidemic spreads and sampling intensity rises. By contrast, high-intensity sampling (*K* = 50) achieves near-certain detection (>95%) with minimal uncertainty. These results demonstrate that early detection is feasible but comes at prohibitively high cost ( USD 100,000 per year for the highest intensity strategies), highlighting the necessity of carefully balancing timing, sampling intensity and site coverage to optimize surveillance performance within realistic budget constraints.

For decision-makers constrained to a total budget of USD 10,000, the optimal strategy identified was *N* = 500 sites with *K* = 10 samples per site. This design achieved a mean detection probability of approximately 75%, although delayed BBTD detection by a year.

## 4. Discussion

Simulation outcomes demonstrate that even moderate, strategically allocated surveillance can substantially improve early detection, enabling rapid response actions—such as roguing or planting material movement restrictions—that reduce epidemic spread and economic losses. By accelerating detection while managing cost, resource-efficient surveillance supports both plant health protection and smallholder resilience.

The reconstructed spatio-temporal patterns suggest that BBTV spread in Benin followed a coherent northward trajectory, with southern departments consistently infected earlier than central and northern regions. The alignment of simulated first-infection timings with independent surveillance data supports the plausibility of the model. These simulations provide a robust foundation for assessing surveillance strategies, highlighting how spatial heterogeneity and epidemic timing influence detection probability. By capturing the broad dynamics of BBTD expansion, the model offers a valuable tool for designing targeted, resource-efficient monitoring and early-detection programs.

Our findings emphasize the critical role of strategic resource allocation in BBTD surveillance particularly in relation to early detection. Detection probability is highly sensitive to both spatial coverage and within-site sampling intensity, with nonlinear interactions governing the speed and certainty of pathogen discovery. While high-intensity sampling across many sites can achieve near-certain detection within a few years, this comes at prohibitively high costs ( USD 100,000 per year). Under realistic budget constraints (e.g., USD 10,000 per year), moderately high coverage with intermediate within-site sampling (*N* = 500, *K* = 10) could still achieve a detection probability of approximately 75% with one year delay, providing a cost-efficient compromise between early warning and expenditure.

The integrated simulation–economic framework presented here quantifies trade-offs between surveillance cost and detection likelihood, providing evidence-based guidance for resource-limited plant health programs. Although this study focused on a cross-sectional survey, the same approach can be extended to longitudinal monitoring, delimitation surveys, and continuous early-warning systems, allowing practitioners to optimize sampling across time and space according to operational constraints and epidemiological objectives. Importantly, although our analysis is grounded in retrospective data from Benin, the insights generated have broader relevance for the design of surveillance strategies in countries with comparable banana production systems where BBTD has not yet been reported. One such example is Ghana, where the endemic presence of the banana aphid vector and the potentially severe impacts on both commercial and smallholder production highlight the critical need for proactive surveillance and control measures [8]. Our assumption in designing surveillance was that the only information available was a general suspicion that a BBTD incursion might have occurred in the country. Consequently, surveillance relied almost exclusively on plant protection agencies to organise BBTD detection activities. A complementary approach is to engage farmers through targeted training in BBTV symptom recognition, coupled with rapid reporting of suspected plants. These elements could be incorporated into the model to enable evaluation of the cost–benefit trade-offs and the value of farmer-assisted BBTD detection programmes.

Recent advances in remote sensing and machine learning for disease detection can be integrated with modelling by providing scalable observational data that can inform and validate surveillance strategies. Retkute et al. [22] demonstrated the potential of combining Landsat-8 satellite data with phenology-informed modelling and machine learning to detect anomalies associated with BBTD at plantation scales; declines in vegetation indices sometimes preceded reported infections by months, illustrating the value of integrating remote sensing for early warning and spatial tracking of disease patterns. Such observational systems could be linked with retrospective simulation frameworks to calibrate model predictions in near real time and refine sampling priorities where on-the-ground surveillance is scarce.

## 5. Conclusions

Optimising pest and disease surveillance under limited resources is critical for protecting farmers’ income and food security in sub-Saharan Africa. This study is the first to quantify both the spatio-temporal spread of BBTV in Benin and the cost–benefit of BBTD surveillance strategies. Early detection is achievable but costly, with high-intensity surveillance nearing certainty at USD 100,000 per year, while moderately high coverage with intermediate sampling delivers high detection at a fraction of the cost, but with anticipated one year delay in detection. Our integrated simulation–economic framework quantifies cost–detection trade-offs and provides transferable guidance for efficient cross-sectional, longitudinal, or delimitation surveys across crops, countries, and pathosystems.

## Supplementary Information

**Figure S1:**
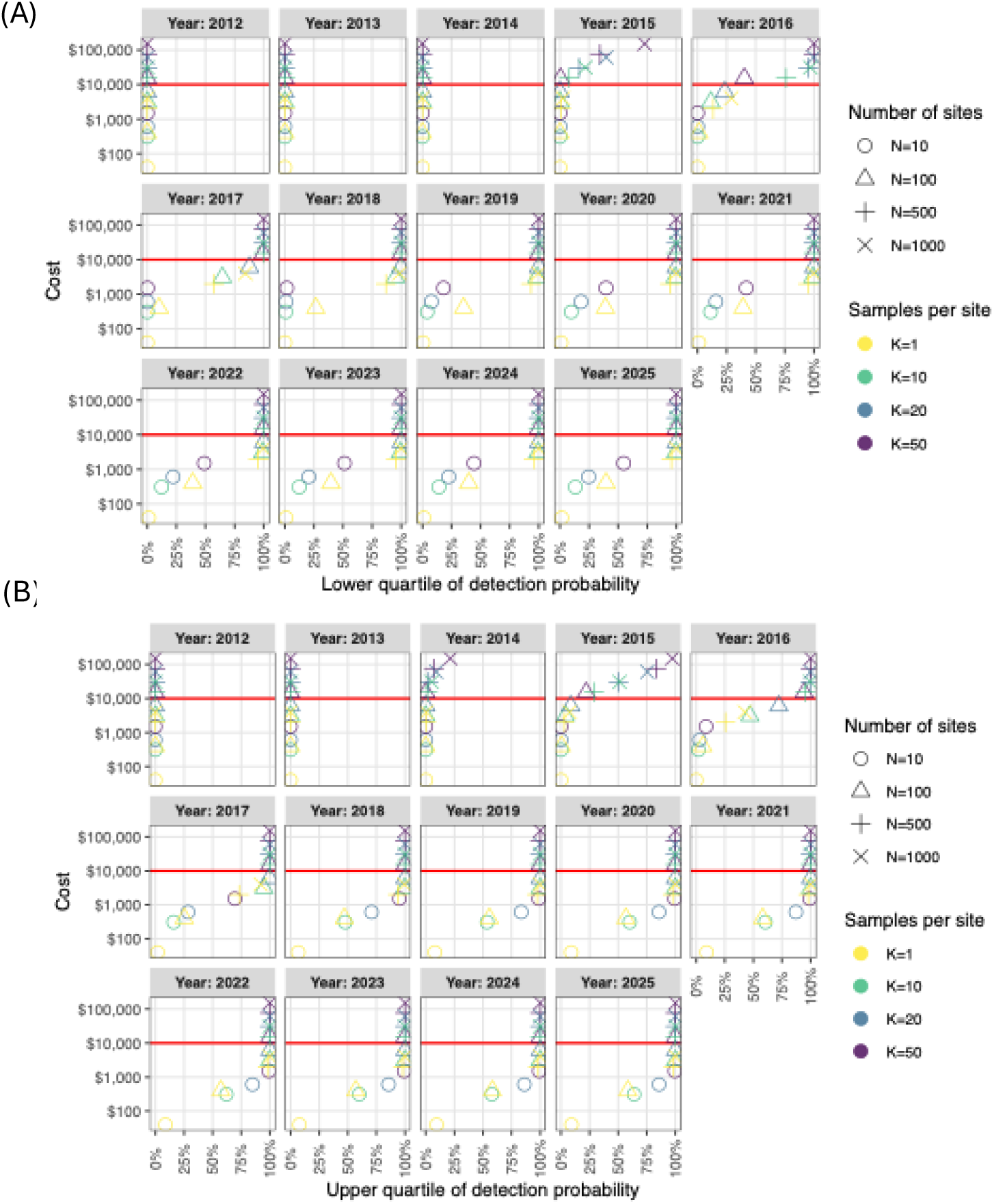
(A) Relationship between lower quartile of detection probability and total annual surveillance cost. (B) Relationship between upper quartile of detection probability and total annual surveillance cost.

